# Genetic analysis of the *STIM1* gene in chronic pancreatitis

**DOI:** 10.1101/691899

**Authors:** Emmanuelle Masson, Wen-Bin Zou, Claudia Ruffert, Vanessa Holste, Patrick Michl, Joachim Mössner, Maren Ewers, Helmut Laumen, Hao Wu, Dai-Zhan Zhou, Zhao-Shen Li, Dong Yu, Arnaud Boulling, Cédric Le Maréchal, David N. Cooper, Jian-Min Chen, Heiko Witt, Jonas Rosendahl, Zhuan Liao, Claude Férec

## Abstract

Chronic pancreatitis is a complex disease that involves many factors, both genetic and environmental. Over the past two decades, molecular genetic analysis of five genes that are highly expressed in human pancreatic acinar cells, namely *PRSS1, PRSS2, SPINK1, CTRC* and *CTRB1*/*CTRB2*, has established that a trypsin-dependent pathway plays a key role in the etiology of chronic pancreatitis. Since Ca^2+^ deregulation can lead to intracellular trypsin activation in experimental acute pancreatitis, we analyzed *STIM1* (encoding stromal interaction molecule-1, the main regulator of Ca^2+^ homeostasis in pancreatic acinar cells) as a candidate modifier gene in French, German and Chinese patients with chronic pancreatitis. The French and German subjects were analyzed by Sanger sequencing whereas the Chinese subjects were analyzed by targeted next-generation sequencing confirmed by Sanger sequencing. A total of 37 rare coding variants (35 missense and 2 nonsense) were identified, which were enriched in patients as compared with controls [2.28% (47/2,057) vs. 0.99% (33/3,322); odds ratio = 2.33, *P* = 0.0001]. This is the first large case-control study to demonstrate a putative association of rare *STIM1* coding variants with chronic pancreatitis. Functional analysis will be required to clarify whether or not the rare *STIM1* variants detected predispose to pancreatitis.

## INTRODUCTION

Chronic pancreatitis is a complex disease that is defined as “a pathologic fibro-inflammatory syndrome of the pancreas in individuals with genetic, environmental and/or other risk factors who develop persistent pathologic responses to parenchymal injury or stress” (Whitcomb et al., 2016). Over the past two decades, molecular genetic analysis of five genes that are highly expressed in human pancreatic acinar cells, namely *PRSS1* encoding cationic trypsinogen (Le Maréchal et al., 2006; Whitcomb et al., 1996), *PRSS2* encoding anionic trypsinogen (Witt et al., 2006), *SPINK1* encoding pancreatic secretory trypsin inhibitor (Witt et al., 2000), *CTRC* encoding chymotrypsin C (Masson et al., 2008; Rosendahl et al., 2008) and *CTRB1-CTRB2* encoding chymotrypsin B1 and B2 (Rosendahl et al., 2018), has established a trypsin-dependent pathway in the etiology of chronic pancreatitis (Hegyi and Sahin-Toth, 2017).

The majority of patients with chronic pancreatitis had prior clinically recognized acute pancreatitis (LaRusch et al., 2015), an acute inflammatory disease of the pancreas postulated to be an autodigestive disease triggered by prematurely activated trypsin within the pancreas (Chiari, 1896). The association of gain-of-function *PRSS1* variants with both recurrent acute pancreatitis and chronic pancreatitis (Gorry et al., 1997; Whitcomb et al., 1996) not only provided support for Chiari’s original hypothesis (Chiari, 1896) but has also contributed to the Sentinel Acute Pancreatitis Event model for the development of chronic pancreatitis (Whitcomb, 1999). Importantly, successive developments of spontaneous acute pancreatitis and chronic pancreatitis have recently been observed in genetically modified mice that carried a heterozygous p.Asp23Ala mutation within the activation peptide of the mouse cationic trypsinogen (Prss1) T7 isoform (the p.Asp23Ala mutant autoactivates to trypsin 50-fold faster than wild-type) (Geisz and Sahin-Tóth, 2018).

The above notwithstanding, our understanding of the early events leading to pancreatitis is still rather limited. In this regard, prolonged and global Ca^2+^ elevation (elicited by bile, alcohol metabolites and other causes) has been described to result in trypsin activation, vacuolization and necrosis of the pancreatic acinar cells in experimental acute pancreatitis (review in (Li et al., 2014)); and stromal interaction molecule-1 (STIM1) is a key regulator for Ca^2+^ homeostasis in both non-excitable and excitable cells (Yuan et al., 2009). These findings suggest that variants in the *STIM1* gene may contribute to the early steps of pancreatitis by disturbing Ca^2+^ homeostasis within the pancreatic tissue.

Variants in the *STIM1* gene have been previously associated a number of diseases such as immunodeficiency and autoimmunity (Picard et al., 2009; Shaw et al., 2013), a novel syndrome of amelogenesis imperfecta and hypohidrosis (Parry et al., 2016), tubular-aggregate myopathy (Bohm et al., 2013; Nesin et al., 2014; Noury et al., 2017), or Stormorken syndrome (Misceo et al., 2014; Morin et al., 2014). Also, tubular aggregate myopathy and Stormorken syndrome patients carrying *STIM1* variants additionally manifested psychiatric disorders (Harris et al., 2017). Moreover, Sofia and colleagues have recently analyzed the *STIM1* gene (included within a panel of 70 genes related to six different pancreatic pathways) in 80 patients with idiopathic chronic pancreatitis (ICP) and found three missense mutations [i.e., c.1310G>A (p.Cys437Tyr), c.1589G>A (p.Arg530His), and c.2246G>A (p.Arg749His)] in different patients (Sofia et al., 2016). In addition to the relatively small number of patients analyzed, this study was limited by the lack of data from a corresponding control population. Herein, we report our findings from a comprehensive variant analysis of the *STIM1* gene in three ICP cohorts.

## PATIENTS AND METHODS

### Patients

This study included 436 French, 517 German and 1,104 Chinese patients with ICP (i.e., absence of both a positive family history and any of the following external precipitating factors, namely alcohol abuse, post-traumatic, hypercalcemic, hyperlipidemic and autoimmune) and corresponding healthy controls. The diagnosis of chronic pancreatitis was made as previously described (Witt et al., 2013; Zou et al., 2016). Informed consent was obtained from each patient and the study was approved by the respective ethics committees.

### Variant screening

The French and German subjects were analyzed by Sanger sequencing; three multiplex PCRs were designed to amplify the entire coding sequence and flanking intronic sequences of the *STIM1* gene (see additional file, Figures. S1 and S2). The Chinese subjects were analyzed by targeted next-generation sequencing followed by Sanger sequencing confirmation, essentially as previously described (Wu et al., 2017; Zou et al., 2018), the primer sequences are provided in Additional file, Figure S3.

### Variant nomenclature and reference sequences

Variant nomenclature followed Human Genome Variation Society recommendations (http://www.hgvs.org/mutnomen/recs.html) (den Dunnen et al., 2016). GenBank accession number NM_003156.3 was used as the *STIM1* mRNA reference sequence. *STIM1* genomic sequence was obtained from human GRCh38/hg38 (https://genome.ucsc.edu/).

### Pathogenicity prediction

This was performed using the Combined Annotation-Dependent Depletion (CADD) method (Kircher et al., 2014) available at https://cadd.gs.washington.edu/.

### Statistical analyses

The assessment of statistical significance of the differences between the carrier frequencies of the *STIM1* variants in patients and controls was performed by the 2×2 contingency table available at http://vassarstats.net/odds2x2.html. The difference was considered as being statistically significant when the *P* value was ≤ 0.05.

## RESULTS AND DISCUSSION

Given the importance of Ca^2+^ signaling for the regulation of pancreatic zymogen activation and the key role of STIM1 in Ca^2+^ homeostasis, we analyzed the *STIM1* gene as a candidate modifier gene for chronic pancreatitis. Employing Sanger sequencing, we first analyzed the entire coding sequence (2,058 bp; NM_003156.3) and exon/intron boundaries of the 12-exon *STIM1* gene in 436 French ICP patients and 1,005 controls, and then repeated this analysis with 517 German ICP patients and 1,121 controls. Our subsequent analysis was limited to coding sequence variants that resulted in amino acid changes and intronic variants that affected canonical donor/acceptor splice sites. Eight such variants were identified in the French cohort and ten in the German cohort; all these variants were single nucleotide substitutions and all were predicted to result in missense substitutions (Additional file, Tables S1 and S2). Since all detected variants were rare variants (defined as having a minor allele frequency of <0.5% in the control population as previously described (Manolio et al., 2009; Tennessen et al., 2012), we first performed aggregate association analysis in the context of each cohort. A significant enrichment of rare variants in patients as compared to controls was noted in the French cohort (odds ratio (OR) = 4.04, *P* = 0.002) but not in the German cohort (OR = 1.64, *P* = 0.26) (Table 1).

**Table 1.** Prevalence of *STIM1* variants in ICP patients versus controls in the French, German and Chinese Populations

| Population | Cases | Controls | Odds ratio | 95% confidence interval | P value |
| --- | --- | --- | --- | --- | --- |
|  | +/n (%) | +/n (%) |  |  |  |
| French | 12/436 (2.75) | 7/1,005 (0.70) | 4.04 | 1.58-10.32 | 0.002 |
| German | 9/517 (1.74) | 12/1,121 (1.07) | 1.64 | 0.69-3.91 | 0.26 |
| Chinese | 26/1,104 (2.36) | 14/1,196 (1.17) | 2.03 | 1.06-3.92 | 0.03 |
| All three combined | 47/2,057 (2.28) | 33/3,322 (0.99) | 2.33 | 1.49-3.65 | 0.0001 |

We also analyzed the *STIM1* gene in 1,104 Chinese ICP patients and 1,196 controls by means of targeted sequencing followed by Sanger sequencing validation. A total of 24 rare variants were identified (Additional file, Table S3), which when taken together were significantly overrepresented in patients as compared to controls (OR = 2.03, *P* = 0.03; Table 1). A Breslow-Day test for homogeneity of the ORs (https://www.prostatservices.com/) between the French, German and Chinese cohorts showed no significant difference (*P* = 0.14). We therefore combined data from these three cohorts (Table 2), the carrier frequency of the aggregated rare variants being significantly higher in patients than in controls (OR = 2.33, *P* = 0.0001; Table 1).

**Table 2.** *STIM1* variants in the combined French, German and Chinese cohorts

| Exon | Nucleotide change | Amino acid change | Patients<br>(*n* = 2,057) |  | Controls<br>(*n* = 3,322) |  | rs number | Allele frequency in gnomAD | CADD score |
| --- | --- | --- | --- | --- | --- | --- | --- | --- | --- |
|  |  |  | + (%) | Population(s) | + (%) | Population(s) |  |  |  |
| Variants detected in patients only |  |  |  |  |  |  |  |  |  |
| 1 | c.91G>C | p.Ala31Pro | 1 (0.05) | F | 0 (0) |  | rs368091975 | 1.19e-5 | 16.10 |
| 1 | c.107C>T | p.Ser36Leu | 1 (0.05) | G | 0 (0) |  | rs200907515 | 3.18e-5 | 13.72 |
| 1 | c.112G>C | p.Ala38Pro | 1 (0.05) | F | 0 (0) |  | rs774499633 | No | 14.95 |
| 1 | c.113C>T | p.Ala38Val | 1 (0.05) | C | 0 (0) |  | No | No | 13.93 |
| 4 | c.454G>A | p.Glu152Lys<sup>a</sup> | 3 (0.15) | C (1), F (2) | 0 (0) |  | rs143916878 | 1.16e-4 | 23.2 |
| 6 | c.747G>C | p.Glu249Asp | 1 (0.05) | C | 0 (0) |  | No | No | 18.57 |
| 9 | c.1231A>G | p.Thr411Ala | 1 (0.09) | C | 0 (0) |  | No | No | 18.54 |
| 11 | c.1498C>T | p.Arg500Trp | 1 (0.05) | C | 0 (0) |  | rs772902514 | 1.59e-5 | 33 |
| 12 | c.1562C>T | p.Ser521Leu | 2 (0.10) | C (1), G (1) | 0 (0) |  | rs745539009 | 1.59e-5 | 24.5 |
| 12 | c.1595G>A | p.Arg532His | 1 (0.05) | C | 0 (0) |  | rs771442242 | 7.96e-6 | 26.5 |
| 12 | c.1615C>T | p.Gln539Ter | 1 (0.05) | C | 0 (0) |  | No | No | 41 |
| 12 | c.1668C>G | p.Ser556Arg | 5 (0.24) | C | 0 (0) |  | rs201543900 | 4.24e-5 | 22.6 |
| 12 | c.1801C>T | p.Pro601Ser | 1 (0.05) | G | 0 (0) |  | rs200960094 | 3.98e-5 | 16.68 |
| 12 | c.1808C>T | p.Ala603Val | 1 (0.05) | C | 0 (0) |  | rs749622475 | 1.19e-5 | 19.32 |
| 12 | c.1843C>T | p.Arg615Cys | 3 (0.15) | C | 0 (0) |  | rs560566339 | 1.19e-5 | 28.5 |
| 12 | c.2012G>A | p.Arg671Gln | 1 (0.05) | C | 0 (0) |  | rs779204802 | 8.04e-6 | 24.3 |
| Variants detected in patients and controls |  |  |  |  |  |  |  |  |  |
| 4 | c.458C>T | p.Thr153Ile | 3 (0.15) | F (1), G (2) | 1 (0.03) | F | rs144602692 | 1.94e-4 | 23.8 |
| 11 | c.1511C>T | p.Thr504Met | 2 (0.10) | C | 1 (0.03) | C | rs146873551 | 8.28e-4 | 20.8 |
| 12 | c.1571C>T | p.Ser524Phe | 5 (0.24) | F (2), G (3) | 8 (0.24) | F (4), G (4) | rs141215990 | 1.78e-3 | 29.1 |
| 12 | c.1589G>A | p.Arg530His | 6 (0.29) | C | 3 (0.09) | C | rs746517083 | 3.18e-5 | 24.5 |
| 12 | c.1612C>T | p.Pro538Ser | 4 (0.19) | F | 1 (0.03) | F | rs35960304 | 5.95e-3 | 20.2 |
| 12 | c.1636G>A | p.Glu546Lys | 2 (0.10) | F (1), G (1) | 2 (0.06) | G | rs371443357 | 3.54e-5 | 23.9 |
| **Variants detected in controls only** |  |  |  |  |  |  |  |  |  |
| 4 | c.408G>C | p.Glu136Asp | 0 (0) |  | 1 (0.03) | C | rs200648767 | 1.77e-4 | 13.79 |
| 4 | c.472C>G | p.Gln158Glu | 0 (0) |  | 1 (0.03) | C | No | No | 21.3 |
| 5 | c.530C>T | p.Thr177Ile | 0 (0) |  | 1 (0.03) | C | rs761973338 | 3.18e-5 | 24.6 |
| 7 | c.826G>C | p.Glu276Gln | 0 (0) |  | 1 (0.03) | C | No | No | 23.0 |
| 8 | c.1010C>T | p.Ser337Phe | 0 (0) |  | 1 (0.03) | G | No | No | 27.6 |
| 11 | c.1499G>A | p.Arg500Gln | 0 (0) |  | 1 (0.03) | C | rs760242778 | 7.97e-6 | 29.4 |
| 11 | c.1505G>A | p.Arg502His | 0 (0) |  | 1 (0.03) | C | rs555016539 | 1.19e-5 | 27.3 |
| 12 | c.1583G>A | p.Ser528Asn | 0 (0) |  | 1 (0.03) | C | rs200078549 | 2.39e-5 | 24.0 |
| 12 | c.1601C>A | p.Ala534Asp | 0 (0) |  | 1 (0.03) | F | No | No | 14.12 |
| 12 | c.1624C>T | p.Arg542Cys | 0 (0) |  | 1 (0.03) | C | rs370846246 | 3.58e-5 | 28.3 |
| 12 | c.1673G>A | p.Arg558Gln | 0 (0) |  | 1 (0.03) | C | rs199503470 | 1.59e-5 | 24.4 |
| 12 | c.1681G>A | p.Glu561Lys | 0 (0) |  | 1 (0.03) | G | rs200557274 | 1.99e-5 | 31 |
| 12 | c.1928G>A | p.Arg643His | 0 (0) |  | 3 (0.09) | G | rs140080199 | 7.57e-4 | 31 |
| 12 | c.1960G>A | p.Ala654Thr | 0 (0) |  | 1 (0.03) | G | rs201466902 | 1.41e-4 | 21.8 |
| 12 | c.2053A>T | p.Lys685Ter | 0 (0) |  | 1 (0.03) | C | No | No | 42 |
| Total |  |  | 47 (2.28) |  | 33 (0.99) |  |  |  |  |

All variants were found in the heterozygous state. C, Chinese. F, French. G, Germany.

Our comprehensive analysis of the *STIM1* gene in three ICP cohorts identified a significant enrichment of rare coding *STIM1* variants in patients as compared to controls by means of aggregate association analysis (Table 1). Notably, none of the identified 37 rare *STIM1* variants correspond to those previously reported to cause or predispose to other diseases (Lacruz and Feske, 2015), potentially strengthening the notion of the tissue-specific effects of different *STIM1* variants. However, pathogenicity prediction by means of the CADD method yielded similar findings among the three groups of variants namely, (i) variants found in only patients, (ii) variants found in both patients and controls and (iii) variants found in only controls (Table 2).

In summary, this is the first large case-control study to demonstrate a putative association of rare *STIM1* coding variants with chronic pancreatitis. Functional analysis will be required to clarify whether or not rare coding *STIM1* variants predispose to pancreatitis.

