## Additional file for "Genetic analysis of the *STIM1* gene in chronic pancreatitis"

**Table S1.** *STIM1* variants identified in the French cohort

| **Exon** | **Nucleotide change** | **Amino acid change** | **Patients (%) (*n* = 436)** | **Controls (%) (*n* = 1,005)** |
|---|---|---|---|---|
| 1 | c.91G>C | p.Ala31Pro | 1 (0.23) | 0 (0) |
| | c.112G>C | p.Ala38Pro | 1 (0.23) | 0 (0) |
| 4 | c.454G>A | p.Glu152Lys | 2 (0.46) | 0 (0) |
| | c.458C>T | p.Thr153Ile | 1 (0.23) | 1 (0.10) |
| 12 | c.1571C>T | p.Ser524Phe | 2 (0.46) | 4 (0.40) |
| | c.1601C>A | p.Ala534Asp | 0 (0) | 1 (0.10) |
| | c.1612C>T | p.Pro538Ser | 4 (0.91) | 1 (0.10) |
| | c.1636G>A | p.Glu546Lys | 1 (0.23) | 0 (0) |
| Total | | | 12 (2.75) | 7 (0.70) |

All variants were found in the heterozygous state.

**Table S2.** *STIM1* variants identified in the German cohort

| **Exon** | **Nucleotide change** | **Amino acid change** | **Patients (%) (*n* = 517)** | **Controls (%) (*n* = 1,121)** |
|---|---|---|---|---|
| 1 | c.107C>T | p.Ser36Leu | 1 (0.19) | 0 (0) |
| 4 | c.458C>T | p.Thr153Ile | 2 (0.38) | 0 (0) |
| 8 | c.1010C>T | p.Ser337Phe | 0 (0) | 1 (0.09) |
| 12 | c.1562C>T | p.Ser521Leu | 1 (0.19) | 0 (0) |
| | c.1571C>T | p.Ser524Phe | 3 (0.58) | 4 (0.36) |
| | c.1636G>A | p.Glu546Lys | 1 (0.19) | 2 (0.18) |
| | c.1681G>A | p.Glu561Lys | 0 (0) | 1 (0.09) |
| | c.1801C>T | p.Pro601Ser | 1 (0.19) | 0 (0) |
| | c.1928G>A | p.Arg643His | 0 (0) | 3 (0.26) |
| | c.1960G>A | p.Ala654Thr | 0 (0) | 1 (0.09) |
| Total | | | 9 (1.74) | 12 (1.07) |

All variants were found in the heterozygous state.

**Table S3.** *STIM1* variants identified in the Chinese cohort

| **Exon** | **Nucleotide change** | **Amino acid change** | **Patients (%) (*n* = 1,104)** | **Controls (%) (*n* = 1,196)** |
|---|---|---|---|---|
| 1 | c.113C>T | p.Ala38Val | 1 (0.09) | 0 (0) |
| 4 | c.408G>C | p.Glu136Asp | 0 (0) | 1 (0.08) |
| | c.454G>A | p.Glu152Lys[a] | 1 (0.09) | 0 (0) |
| | c.472C>G | p.Gln158Glu | 0 (0) | 1 (0.08) |
| 5 | c.530C>T | p.Thr177Ile | 0 (0) | 1 (0.08) |
| 6 | c.747G>C | p.Glu249Asp | 1 (0.09) | 0 (0) |
| 7 | c.826G>C | p.Glu276Gln | 0 (0) | 1 (0.08) |
| 9 | c.1231A>G | p.Thr411Ala | 1 (0.09) | 0 (0) |
| 11 | c.1498C>T | p.Arg500Trp | 1 (0.09) | 0 (0) |
| | c.1499G>A | p.Arg500Gln | 0 (0) | 1 (0.08) |
| | c.1505G>A | p.Arg502His | 0 (0) | 1 (0.08) |
| | c.1511C>T | p.Thr504Met | 2 (0.18) | 1 (0.08) |
| 12 | c.1562C>T | p.Ser521Leu[b] | 1 (0.09) | 0 (0) |
| | c.1583G>A | p.Ser528Asn | 0 (0) | 1 (0.08) |
| | c.1589G>A | p.Arg530His | 6 (0.54) | 3 (0.25) |
| | c.1595G>A | p.Arg532His | 1 (0.09) | 0 (0) |
| | c.1615C>T | p.Gln539Ter | 1 (0.09) | 0 (0) |
| | c.1624C>T | p.Arg542Cys | 0 (0) | 1 (0.08) |
| | c.1668C>G | p.Ser556Arg | 5 (0.45) | 0 (0) |
| | c.1673G>A | p.Arg558Gln | 0 (0) | 1 (0.08) |
| | c.1808C>T | p.Ala603Val | 1 (0.09) | 0 (0) |
| | c.1843C>T | p.Arg615Cys | 3 (0.27) | 0 (0) |
| | c.2012G>A | p.Arg671Gln | 1 (0.09) | 0 (0) |
| | c.2053A>T | p.Lys685Ter | 0 (0) | 1 (0.08) |
| Total | | | 26 (2.36) | 14 (1.17) |

Variants were not found in the French and German cohorts unless otherwise stated. All variants were found in the heterozygous state.

[a]Also found in two French patients.

[b]Also found in one German patient.

**A**

Primers employed for both amplification and sequencing (5'>3')

| | |
|---|---|
| STIM1_exon1F | AAGCTGGGACTTGATCCTTTG |
| STIM1_exon1R | CATGTACAAACCTAGATTTCAACTTGC |
| STIM1_exon2F | GCCAGTTAAGTGCCTACATGA |
| STIM1_exon2R | TCCCTGATAGGGATGAGTTCCT |
| STIM1_exon3F | GAATGTGTTATGGCTAGCTAGAG |
| STIM1_exon3R | ATGTTCTTCTAAGGCCAAGTTGC |
| STIM1_exon4F | GTGGTAAATATTAAGGTCAGCATGAC |
| STIM1_exon4R | TAACTGGCCAGAGCAATCTG |
| STIM1_exon5F | CTCCATATACCCTGTTGAGACCCA |
| STIM1_exon5R | TGCTATCTTGCAGGTGTGCCTA |
| STIM1_exon6F | TCTGTTATGGAAGGCTTCATAGAG |
| STIM1_exon6R | AGTTTTGGAAGGATGATGCAGC |
| STIM1_exon7F | AGCTGTCATTTTCCTCTTTGATGC |
| STIM1_exon7R | TGACTCTAGAACATAGTCTTTGGATC |
| STIM1_exon8-10F | TCTCCTTCCTGGGAGAGTTGT |
| STIM1_exon8-10R | ACTATCTTCCTGGGTATCTGCC |
| STIM1_exon11F | TCATGGGCACCTCCTTACCT |
| STIM1_exon11R | AGACTGTCCAGATGAAAGGCTG |
| STIM1_exon12F | TCCTTGTCTTCTCGTGTTGTC |
| STIM1_exon12R | ACAGCAACTAAGACATGCACTG |

Primer employed for sequencing exon 9

| | |
|---|---|
| STIM1_exon9_seqR | ATGGAATTGGTGTTGGAG |

**B**

Exon 1

```
>chr11:3855429-3856502 1074bp
AAGCTGGGACTTGATCCTTTGcgcgggatcctggcaaagactagcgcgggccgggggtccgggagagcccgctag
gggcggggattccggggagccgtcttcaccggttattccgggatccagctgggcgctggggctggcccgggcttc
gctggggaccgggcggcgcggggcgggcgcggagacgcacgcccccgcccgccccgggcccgccccgcgccgccc
gcccgcctggaagccgctgtcctgggcctggccggtgtgcgtccgcctgctggacctgggcaccgccagccgcct
gggcacgggactgggcgggggcgctgacctcggcctaggaggcccaggatcccggagacgcccgcgccctcagga
ccctgcgggtcgcacgccctccccagcttctgctgctcgccgctcttcggcagggcgaggtcaggtgccccccttc
tcgcctctcttctcttctcttctcttcctcctccacttctgtgcccgcggagactccggccgccccccttccgcag
gggtgtagtaatctgcggagctgacagcagccccgcagccaccctgcccgaagtctccggaagcggcacgagctc
aggccgccgcagccccggcggacccactgttggacctgaggagccagccctcctcccgcacccaaacttggagca
cttgacctttggctgttggagggggcaggctcgcgggtggctggacagctgcggagccgcgagggcatcttgcct
ggagaccgtcggctgcactcccgggctcctggctttgcctctgggatcccgaggtgtccacatcagacgcatgtt
gactgagacctagagtcatggatgtatgcgtccgtcttgccctgtggctcctctggggactcctcctgcaccagg
gccagagcctcagccatagtcacagtgagaaggcgacaggaaccagctcgggggccaactctgaggagtccactg
cagcaggtaaggccttgctgcgggctggactgggctggaggctttggctcaggactgagtggcccgaagtggGCA
AGTTGAAATCTAGGTTTGTACATG
```

***(to be continued)***

**Figure S1.** PCR primers used to analyze the *STIM1* gene by Sanger sequencing in the French and German cohorts. (**A**) Sequences of the primers. F, forward. R, reverse. (**B**) Positions of the primers (in capital letters) in the context of the *STIM1* genomic sequence (hg38). The coding sequence of the *STIM1* gene (NM_003156.3) is highlighted in blue. The translational initiation and termination codons are boxed. The primer employed to sequence exon 9 is highlighted in red.

### Exon 2

>chr11:3967434-3967818 385bp
GCCAGTTAAGTGCCTACATGAggagtcagatgctagaagctaaggatgctgacacaggtgggtgataggttctgg
gtggcagctctgagtaattttgtctcttgcttttcttacacagagttttgccgaattgacaagcccctgtgtcac
agtgaggatgagaaactcagcttcgaggcagtccgtaacatccacaaactgatggacgatgatgccaatggtgat
gtggatgtggaagaaagtgatgaggtgagctctcccatcctgctatgtctcttttcttcctgtgtgaattagt
ctttgaaagctacttagacagcctcttccctggggagagatgttctctgctggcttgttctttAGGAACTCATCC
CTATCAGGGA

### Exon 3

>chr11:4023774-4024069 296bp
GAATGTGTTATGGCTAGCTAGAGgcaggtgacctgtgtggagatcttgacgggctgactcctaggtatctcttgt
gacttgtgtacttttcccttgcagttcctgagggaagacctcaattaccatgacccaacagtgaaacacagcacc
ttccatggtgaggataagctcatcagcgtggaggacctgtggaaggcatggaagtcatcagaaggtaataggcag
cctggtcatcaatcctagttgtgggaaggtttagagaagagagaagcaGCAACTTGGCCTTAGAAGAACAT

### Exon 4

>chr11:4055440-4055722 283bp
GTGGTAAATATTAAGGTCAGCATGACaacaaatgaaagcagtgcttggcattctagagtcatggctttgcttgtc
tcttttcacagtatacaattggaccgtggatgaggtggtacagtggctgatcacatatgtggagctgcctcagta
tgaggagaccttccggaagctgcagctcagtggccatgccatgccaaggtcaggaggggactgggtttttctctg
ttgagggtacggggaatgggctggagtgggcctgccttCAGATTGCTCTGGCCAGTTA

### Exon 5

>chr11:4059113-4059507 395bp
CTCCATATACCCTGTTGAGACCCAactggtatagactctgtgttctagagtgcagaggggagcaatcaccaagag
ctagaagtgttcctggggggaggcgggtaatcctaccaggatccttcctggctttactgggagggaactgatctgc
tactctttgcctcaacaggctggctgtcaccaacaccaccatgacagggactgtgctgaagatgacagaccggag
tcatcggcagaagctgcagctgaaggctctggatacagtgctctttgggcctcctctctgtgagtcttgtgttga
gaagggctactgctgtgccatggaaaccaaagctgttgtagtggataggctaaggctaaggctctgggtctccTA
GGCACACCTGCAAGATAGCA

### Exon 6

>chr11:4069926-4070338 413bp
TCTGTTATGGAAGGCTTCATAGAGgagggatgcagtggagtctgcaaggctaagtgtgcagtgggcaccctaact
catcatgccctcccctctctggcagtgactcgccataatcacctcaaggacttcatgctggtggtgtctatcgtt
attggtgtgggcggctgctggtttgcctatatccagaaccgttactccaaggagcacatgaagaagatgatgaag
gacttggaggggttacaccgagctgagcagagtctgcatgaccttcaggaaaggtaaggcctgccccttcaggaa
aggtgaggccctgccagttcttgggaacctcctatttccacctgggtgtgagccacttggcccctgagaccttgt
cagcatggcagcccagGCTGCATCATCCTTCCAAAACT

### Exon 7

>chr11:4074408-4074805 398bp
AGCTGTCATTTTCCTCTTTGATGCcatgactcatggcatgttggctggcacccccttgcctggcctcctccagct
ccctgcattgccccccagg ctgcacaaggcccaggaggagcaccgcacagtggaggtggagaaggtccatctgg
aaaagaagctgcgcgatgagatcaaccttgctaagcaggaagcccagcggctgaaggagctgcgggagggtactg
agaatgagcggagccgccaaaaatatgctgaggaggagttggagcaggtaggagagtccacaaattcctggacac
cttgacgggtgggtaaagggcaggggcccagggtctggctagaaagttacatgtgtgaggaatttgaaatatGAT
CCAAAGACTATGTTCTAGAGTCA

### Exons 8 to 10

>chr11:4082068-4083611 1544bp
TCTCCTTCCTGGGAGAGTTGTaaagcagataagaagtctgagttctgaagcatcatacagagatgttggagagtc
ttagtagcagtaaatgaactcacatccttttggctgcctaggttcgggaggccttgaggaaagcagagaaggagc
tagaatctcacagctcatggtatgctccagaggcccttcagaagtggctgcagctgacacatgaggtggaggtgc
aatattacaacatcaagaagcaaaatgctgagaagcagctgctggtggccaaggaggggtgagaacagcccttc
tattgtcctcttttctcctttttgcccttctcctttttacctgcatatcttcctcttctctccctctccctgttc
ctcccattgggtgaagagatatcctggtggtgggttctgtttctttcctaatgttcagctcctgacagtgtcctc

**Figure S1 *(continued)***

aggcttctggaggagggaatgagggtgtaggggcattcctgcctgtcttctgggaaagatgaaacaagtggaggc
aggcagagggaaagctctgccagccccaggctgccatggggacagaggtcctcctggggcaaaagggctatcaag
gacccctttgcttctcccctcgctttctctatttgtcccctcctttttcttctaccttgcctgcctttctctt
tttcctctttctcttccctttccttgtacagcctcagttgtgctaggaggagtgggccccagccataggggaagg
cctttctcatttattccattctcgaatccctgctcttttttgagctgggggcctcatctttgcaggctgagaagat
aaaaaagaagagaaacacactctttggcaccttccacgtggcccacagctcttccctggatgatgtagatcataa
aattctaacagctaagtaagtaacaccagttatctactctggcaatgtccatatcatcctcagaatttgagggca
tacagggcCTCCAACACCAATTCCATttcctcattggtggagggacagctctggtctcctgcctcagcctttatt
tatctcctgccttgctcttaagtttgagtttattgtgtgttttattcacacatattctcaaaacttgttcctctg
agaagaggcttcattcctattggggctcacaccaagtccatgcctgcagttctctttcctctgtcttcaggcaag
cactgagcgaggtgacagcagcattgcgggagcgcctgcaccgctggcaacagatcgagatcctctgtggcttcc
agattgtcaacaaccctggcatccactcactggtggctgccctcaacatagaccccagctggatgggcagtacac
gccccaaccctgctcacttcatcatgactgacgacgtggatgacatggatgaggagattgtgtctcccttgtcca
tgcagtgtaggtgacctctttgcggggatgaaggaaggagcctttgatgtacaggttgagaagtctgtggcctct
gttgtgcttaaacagcagatggGGCAGATACCCAGGAAGATAGT

Exon 11
>chr11:4086404-4086625 222bp
TCATGGGCACCTCCTTACCTgccagcccaaagtgggctggcccctcctgacactttctttattctccttgcagcc
cctagcctgcagagcagtgttcggcagcgcctgacggagccacagcatggcctgggatctcagaggttggtagag
ggcgaggctggccacttcttgacaagccgggtatctctgcggcgaatgcgCAGCCTTTCATCTGGACAGTCT

Exon 12
>chr11:4091183+4092071 889bp
TCCTTGTCTTCTCGTGTTGTCcctctctcctcttcgcctttcccctatcacctcatccaatatatgtccctttct
tcctctctgccccatgtcttgcagggatttgacccattccgattcggagtcctccctccacatgagtgaccgcca
gcgtgtggcccccaaacctcctcagatgagccgtgctgcagacgaggctctcaatgccatgacttccaatggcag
ccaccggctgatcgagggggtccacccagggtctctggtggagaaactgcctgacagccctgccctggccaagaa
ggcattactggcgctgaaccatgggctggacaaggcccacagcctgatggagctgagcccctcagccccacctgg
tggctctccacatttggattcttcccgttctcacagccccagctccccagacccagacacaccatctccagttgg
ggacagccgagccctgcaagccagccgaaacacacgcattccccacctggctggcaagaaggctgtggctgagga
ggataatggctctattggcgaggaaacagactccagcccaggccggaagaagtttcccctcaaaatctttaagaa
gcctcttaagaagtaggcaggatggggtggcagtaaagggacagcttgtccttccctgggtgttctgtctctcct
tccctcccttccttcaagataactggccccaagagtggggcatgggaagggctggtccaggggtctgggcactgt
acatacctgccccctcatccttgggtccttcattattatttattaactgaccaccatggcctgcctgccctgcct
ccgtcccaaccatgggctgctgctgtcactccctctccacttCAGTGCATGTCTTAGTTGCTGT

**Figure S1 *(continued)***

**PCR mix 1**

| | |
|---|---|
| STIM1_exon2F (10 µM) | 0.5 µL |
| STIM1_exon2R (10 µM) | 0.5 µL |
| STIM1_exon3F (10 µM) | 0.5 µL |
| STIM1_exon3R (10 µM) | 0.5 µL |
| STIM1_exon4F (10 µM) | 0.5 µL |
| STIM1_exon4R (10 µM) | 0.5 µL |
| STIM1_exon5F (10 µM) | 0.5 µL |
| STIM1_exon5R (10 µM) | 0.5 µL |
| STIM1_exon6F (10 µM) | 0.5 µL |
| STIM1_exon6R (10 µM) | 0.5 µL |
| STIM1_exon7F (10 µM) | 0.5 µL |
| STIM1_exon7R (10 µM) | 0.5 µL |
| STIM1_exon12F (10 µM) | 0.5 µL |
| STIM1_exon12R (10 µM) | 0.5 µL |
| Qiagen Multiplex PCR Kit (2×) | 12.5 µL |
| DNA (50 ng/µL) | 1.0 µL |
| Water | 4.5 µL |
| Total | 25 µL |

PCR program:
Initial denaturation at 95°C for 15 min, followed by 40 cycles of denaturation at 95°C for 30 secs, annealing at 62.5°C for 30 secs and extension at 72°C for 2min, with a final elongation step at 72°C for 5 min.

**PCR mix 2**

| | |
|---|---|
| STIM1_exon8-10F | 0.2 µL |
| STIM1_exon8-10R | 0.2 µL |
| Qiagen Multiplex PCR Kit (2×) | 5 µL |
| DNA (50 ng/µL) | 1 µL |
| Water | 3.6 µL |
| Total | 10 µL |

PCR program:
Initial denaturation at 95°C for 15 min, followed by 35 cycles of denaturation at 95°C for 30 secs, annealing at 62.5°C for 30 secs and extension at 72°C for 2min, with a final elongation step at 72°C for 5min.

**PCR mix 3**

| | |
|---|---|
| STIM1_exon1F | 0.2 µL |
| STIM1_exon1R | 0.2 µL |
| Qiagen Multiplex PCR Kit (2×) | 5 µL |
| Q solution Qiagen | 2 µL |
| DNA (50 ng/µL) | 1.0 µL |
| Water | 1.6 µL |
| Total | 10 µL |

PCR program:
Initial denaturation at 95°C for 15 min, followed by 35 cycles of denaturation at 95°C for 30 secs, annealing at 61°C for 30 secs and extension at 72°C for 90 secs, with a final elongation step at 72°C for 5 min.

**Figure S2.** Multiplex PCR mixes and conditions used to analyze the *STIM1* gene by Sanger sequencing in the French and German cohorts. Qiagen Multiplex PCR Kit, Cat. 206145 (Courtaboeuf, France).

#### Exon 1

Exon1_F: CCACATCAGACGCATGTTGACT
Exon1_R: AACCTAGATTTCAACTTGCCCACTT
>chr11:3856236-3856494 259bp
CCACATCAGACGCATGTTGACTgagacctagagtcatggatgtatgcgtccgtcttgccctgtggctcctctggg
gactcctcctgcaccagggccagagcctcagccatagtcacagtgagaaggcgacaggaaccagctcgggggcca
actctgaggagtccactgcagcaggtaaggccttgctgcgggctggactgggctggaggctttggctcaggactg
agtggcccgAAGTGGGCAAGTTGAAATCTAGGTT

#### Exon 2

Exon2_F: TGACACAGGTGGGTGATAGGTT
Exon2_R: GGCTGTCTAAGTAGCTTTCAAAGACTA
>chr11:3967483-3967756 274bp
TGACACAGGTGGGTGATAGGTTctgggtggcagctctgagtaattttgtctcttgcttttcttacacagagtttt
gccgaattgacaagcccctgtgtcacagtgaggatgagaaactcagcttcgaggcagtccgtaacatccacaaac
tgatggacgatgatgccaatggtgatgtggatgtggaagaaagtgatgaggtgagctctcccatcctgctatgtc
tctcttttcttcctgtgtgaatTAGTCTTTGAAAGCTACTTAGACAGCC

#### Exon 3

Exon3_F: AGGTGACCTGTGTGGAGATCTT
Exon3_R: ATGTTCTTCTAAGGCCAAGTTGCT
>chr11:4023799-4024069 271bp
AGGTGACCTGTGTGGAGATCTTgacgggctgactcctaggtatctcttgtgacttgtgtacttttcccttgcagt
tcctgagggaagacctcaattaccatgacccaacagtgaaacacagcaccttccatggtgaggataagctcatca
gcgtggaggacctgtggaaggcatggaagtcatcagaaggtaataggcagcctggtcatcaatcctagttgtggg
aaggtttagagaagagagaagcAGCAACTTGGCCTTAGAAGAACAT

#### Exon 4

Exon4_F: GGTCAGCATGACAACAAATGAAAGC
Exon4_R: TAACTGGCCAGAGCAATCTGAAG
>chr11:4055454-4055722 269bp
GGTCAGCATGACAACAAATGAAAGCagtgcttggcattctagagtcatggctttgcttgtctcttttcacagtat
acaattggaccgtggatgaggtggtacagtggctgatcacatatgtggagctgcctcagtatgaggagaccttcc
ggaagctgcagctcagtggccatgccatgccaaggtcaggaggggactgggtttttctctgttgagggtacgggg
aatgggctggagtgggcctgcCTTCAGATTGCTCTGGCCAGTTA

#### Exon 5

Exon5_F: CGGGTAATCCTACCAGGATCCTT
Exon5_R: GCCTTAGCCTATCCACTACAACAG
>chr11:4059210-4059468 259bp
CGGGTAATCCTACCAGGATCCTTcctggctttactgggagggaactgatctgctactctttgcctcaacaggctg
gctgtcaccaacaccaccatgacagggactgtgctgaagatgacagaccggagtcatcggcagaagctgcagctg
aaggctctggatacagtgctctttggcctcctctctgtgagtcttgtgttgagaagggctactgctgtgccatg
gaaaccaaagCTGTTGTAGTGGATAGGCTAAGGC

***(to be continued)***

**Figure S3.** PCR primers used for the analysis of the *STIM1* gene by targeted next generation sequencing in the Chinese cohort. Positions of the primers (in capital letters) are annotated in the context of the *STIM1* genomic sequence (hg38). The coding sequence of the *STIM1* gene (NM_003156.3) is highlighted in blue. The translational initiation and termination codons are boxed. Different primer pairs in the same exon are distinguished by different underlining styles.

### Exon 6

Exon6_1_F: GCCTCTCATTTCAAAACAAAGCCT
Exon6_1_R: CAATAACGATAGACACCACCAGCAT
Exon6_2_F: GCAGTGACTCGCCATAATCACC
Exon6_2_R: GGAAATAGGAGGTTCCCAAGAACTG
>chr11:4069805-4070264 460bp
GCCTCTCATTTCAAAACAAAGCCTtggctagcagagagatagacatagagcttactgtaatagtgtatggcaagt
gtgtatctaagaagagaaactaattccttctcagtgggagagtgtgtctgttatggaaggcttcatagaggaggg
atgcagtggagtctgcaaggctaagtgtgcagtgggcaccctaactcatcatgccctcccctctctgGCAGTGAC
TCGCCATAATCACCtcaaggacttcATGCTGGTGGTGTCTATCGTTATTGgtgtgggcggctgctggtttgccta
tatccagaaccgttactccaaggagcacatgaagaagatgatgaaggacttggaggggttacaccgagctgagca
gagtctgcatgaccttcaggaaaggtaaggcctgccccttcaggaaaggtgaggccctgcCAGTTCTTGGGAACC
TCCTATTTCC

### Exon 7

Exon7_1_F: GGTTTCTATGGGCCTTGAGCTA
Exon7_1_R: TGCTTAGCAAGGTTGATCTCATCG
Exon7_2_F: GGTGGAGAAGGTCCATCTGGAA
Exon7_2_R: TCAAGGTGTCCAGGAATTTGTGG
>chr11:4074335-4074712 378bp
GGTTTCTATGGGCCTTGAGCTAgctcagagccagacacacagcagagcagggagaatatatgctgagagttggag
ctgtcattttcctctttgatgccatgactcatggcatgttggctggcaccccttgcctggcctcctccagctcc
ctgcattgccccccagctgcacaaggcccaggaggagcaccgcacagtggaGGTGGAGAAGGTCCATCTGGAA
aagaagctgcgCGATGAGATCAACCTTGCTAAGCAggaagcccagcggctgaaggagctgcgggagggtactgag
aatgagcggagccgccaaaaatatgctgaggaggagttggagcaggtaggagagtCCACAAATTCCTGGACACCT
TGA

### Exon 8

Exon8_F: CTTAGTAGCAGTAAATGAACTCACATCCT
Exon8_R: TATGCAGGTAAAAAGGAGAAGGGCAAAAAG
>chr11:4082142-4082414 273bp
CTTAGTAGCAGTAAATGAACTCACATCCTtttggctgcctaggttcgggaggccttgaggaaagcagagaaggag
ctagaatctcacagctcatggtatgctccagaggcccttcagaagtggctgcagctgacacatgaggtggaggtg
caatattacaacatcaagaagcaaaatgctgagaagcagctgctggtggccaaggaggggtgagaacagccctt
ctattgtcctcttttctcCTTTTTGCCCTTCTCCTTTTTACCTGCATA

### Exon 9

Exon9_F: CCATTCTCGAATCCCTGCTCTT
Exon9_R: CCTCAAATTCTGAGGATGATATGGACATTG
>chr11:4082833-4083039 207bp
CCATTCTCGAATCCCTGCTCTTtttgagctgggggcctcatctttgcaggctgagaagataaaaaagaagagaaa
cacactctttggcaccttccacgtggcccacagctcttccctggatgatgtagatcataaaattctaacagctaa
gtaagtaacaccagttatctactctggCAATGTCCATATCATCCTCAGAATTTGAGG

### Exon 10

Exon10_1_F: GCCTTGCTCTTAAGTTTGAGTTTATTGT
Exon10_1_R: GGATGCCAGGGTTGTTGACAAT
Exon10_2_F: AGATCGAGATCCTCTGTGGCTT
Exon10_2_R: ACATCAAAGGCTCCTTCCTTCATC
>chr11:4083126-4083542 417bp
GCCTTGCTCTTAAGTTTGAGTTTATTGTgtgttttattcacacatattctcaaaacttgttcctctgagaagagg
cttcattcctattggggctcacaccaagtccatgcctgcagttctctttcctctgtcttcaggcaagcactgagc
gaggtgacagcagcattgcgggagcgcctgcaccgctggcaacAGATCGAGATCCTCTGTGGCTTccagATTGTC
AACAACCCTGGCATCCactcactggtggctgccctcaacatagaccccagctggatgggcagtacacgccccaac
cctgctcacttcatcatgactgacgacgtggatgacatggatgaggagattgtgtctcccttgtccatgcagtgt
aggtgacctctttgcgggGATGAAGGAAGGAGCCTTTGATGT

**Figure S3 *(continued)***

### Exon 11

```
Exon11_F: GGTGGTCTCCAGCAAGCATTTA
Exon11_R: GAAAGACTGTCCAGATGAAAGGCT
>chr11:4086381-4086628 248bp
GGTGGTCTCCAGCAAGCATTTAttcatgggcacctccttacctgccagcccaaagtgggctggcccctcctgaca
ctttctttattctccttgcagcccctagcctgcagagcagtgttcggcagcgcctgacggagccacagcatggcc
tgggatctcagaggttggtagagggcgaggctggccacttcttgacaagccgggtatctctgcggcgaatgcgcA
GCCTTTCATCTGGACAGTCTTTC
```

### Exon 12

```
Exon12_1_F: CCCTATCACCTCATCCAATATATGTCC
Exon12_1_R: GCGCCAGTAATGCCTTCTTG
Exon12_2_F: GACGAGGCTCTCAATGCCAT
Exon12_2_R: CCAACTGGAGATGGTGTGTCT
Exon12_3_F: ATTTGGATTCTTCCCGTTCTCACA
Exon12_3_R: AAGGACAAGCTGTCCCTTTACTG
>chr11:4091225-4091835 611bp
CCCTATCACCTCATCCAATATATGTCCctttcttcctctctgccccatgtcttgcagggatttgacccattccga
ttcggagtcctccctccacatgagtgaccgccagcgtgtggcccccaaacctcctcagatgagccgtgctgcaGA
CGAGGCTCTCAATGCCATgacttccaatggcagccaccggctgatcgaggggtccacccagggtctctggtgga
gaaactgcctgacagccctgccctggcCAAGAAGGCATTACTGGCGCtgaaccatgggctggacaaggcccacag
cctgatggagctgagcccctcagccccacctggtggctctccacATTTGGATTCTTCCCGTTCTCACAgccccag
ctccccagacccAGACACACCATCTCCAGTTGGggacagccgagccctgcaagccagccgaaacacacgcattcc
ccacctggctggcaagaaggctgtggctgaggaggataatggctctattggcgaggaaacagactccagcccagg
ccggaagaagtttcccctcaaaatctttaagaagcctcttaagaagtaggcaggatggggtggCAGTAAAGGGAC
AGCTTGTCCTT
```

**Figure S3 *(continued)***
